# Macroscale coupling between structural and effective connectivity in the mouse brain

**DOI:** 10.1101/2023.02.22.529400

**Authors:** Danilo Benozzo, Giorgia Baron, Ludovico Coletta, Alessandro Chiuso, Alessandro Gozzi, Alessandra Bertoldo

## Abstract

How the emergent functional connectivity (FC) relates to the underlying anatomy (structural connectivity, SC) is one of the biggest questions of modern neuroscience. At the macro-scale level, no one-to-one correspondence between structural and functional links seems to exist. And we posit that to better understand their coupling, two key aspects should be taken into account: the directionality of the structural connectome and the limitations of describing network functions in terms of FC. Here, we employed an accurate directed SC of the mouse brain obtained by means of viral tracers, and related it with single-subject effective connectivity (EC) matrices computed by applying a recently developed DCM to whole-brain resting-state fMRI data. We analyzed how SC deviates from EC and quantified their couplings by conditioning both on the strongest SC links and EC links. We found that when conditioning on the strongest EC links, the obtained coupling follows the unimodal-transmodal functional hierarchy. Whereas the reverse is not true, as there are strong SC links within high-order cortical areas with no corresponding strong EC links. This mismatch is even more clear across networks. Only the connections within sensory motor networks align both in terms of effective and structural strength.

## Introduction

How structural connectivity (SC) is coupled with functional brain properties remains an open question for modern neuroscience (Suárez et al., 2020). This problem heavily depends on the scale used to investigate it. Even at the micro scale, where a detailed microcircuit structure can be coupled with biophysical models, there is still discordance in reproducing empirical functional properties (Lynn and Bassett, 2019). This is more pronounced in larger scales, when bottom-up models become intractable due to their fast complexity growth and top-down models are the solely alternative. Focusing on the macro scale case, which is the target of this work, the top-down models currently available are of two main types (Li and Yap, 2022; D’Angelo and Jirsa, 2022): brain network model (BNM) (Ritter et al., 2013) and dynamic causal modeling (DCM) (Friston et al., 2003). In both cases, SC is often used to constrain the interactions across brain units. In BNM, this is a strong a priori assumption, because single unit brain dynamics are placed on top of a structural matrix. Within the DCM framework, even if in principle there is no need for structurally-informed prior, several variants have been developed with structural information (Crimi et al., 2021) with the aim of both reducing the dimensionality of the model and to meet the assumption that a functionally effective link implies an underlying structural link.

However, given that from micro to macro scale functional connectivity (FC) deviates more and more from SC (Miŝic et al., 2016), there is no clear consensus that a structurally-informed prior is always beneficial. A possible explanation for this phenomenon could be that the contribution of emergent network properties overcome the importance of a single brain region, conflicting with the pairwise nature of the structural information. Further works also investigated the heterogeneity in the structural-functional coupling across neocortical areas, and found that the overlapping between FC and SC is maximal in primary sensory and motor regions, and it gradually decreases toward a global minimum in transmodal brain areas (Atasoy et al., 2016; Vázquez-Rodríguez et al., 2019; Preti and Ville, 2019; Liu et al., 2022a). This latter result is in line with the previous work showing that a sensory fugal gradient in the spatial organization of FC represents one fundamental axis capturing the intrinsic architecture of the cortex (Margulies et al., 2016), thus supporting the high degree of complexity in the relation between anatomical connections and functional properties of the brain.

A more recent line of research has focused on how SC relates with the dynamics of brain activity, thus going beyond the static steady-state nature of FC. For example, Liu et al. (2022b) studied how the coupling between SC and dynamic FC evolves in time, while Gu et al. (2022) tried to relate functional modular flexibility with structure by means of a measure of control theory. Lastly, Avena-Koenigsberger et al. (2017) provided a comprehensive overview of key aspects of communication dynamics and their link with topological properties of SC. This led us to the effective connectivity (EC) matrix as a proxy to study brain dynamics. Traditionally, EC refers to the causal influence that each element of a system exerts on the dynamics of the other elements (Bullmore and Sporns, 2009). EC is not directly measured, whereas it is defined in the context of DCM and derived by a process of model inversion. Indeed, the main limitation of DCM was the scalability to whole-brain analysis. This has been recently faced in Razi et al. (2017); Frassle et al. (2021); Prando et al. (2020).

Here, our interest is in trying to understand whether and how SC and EC relate, which is still unclear. However, human EC models lack intrinsic validation, owing to unavailability of reliable information regarding the directed anatomical connectivity of the human brain. Indeed, in human studies, SC is commonly reconstructed from diffusion-weighted MRI which is non-invasive and easily available, but it is not accurate in the tract reconstruction (Schirner et al., 2022; Maier-Hein et al., 2017). Moreover, it provides a symmetric connectivity matrix, a factor that clashes with EC being asymmetric (Kale et al., 2018). On the other hand, viral tracer techniques on animal models provide an accurate reconstruction of monosynaptic axonal path and are considered as the gold standard for mapping the structural connectome (Grandjean et al., 2017, Coletta et al., 2020). Regarding the mouse brain, its mesoscale connectome has been mapped via directional viral traces (Oh et al., 2014). This resource represents the ideal framework for studying the coupling between structural connectome and effective connectivity because of its directed organization and achieved resolution, which surpasses what is doable with MRI in primates and humans.

Here, we studied the relation between effective connectivity and directed structural connectivity at the whole-brain level in the mouse brain. EC was computed subject-wise on a dataset of resting-state fMRI BOLD signals recorded from 20 anesthetized mice by means of sparse DCM (Prando et al., 2020). Global directed SC was inferred from a directed weighted voxel-wise model of the mouse brain (Knox et al., 2018) obtained through viral tracings. At the global level, we showed that the strongest structural links are associated with the strongest EC links, and vice-versa. However, a more detailed analysis at the node level revealed that the coupling strength changes in a network dependent fashion based on whether the link selection was based on the strongest EC or SC links. Moreover, we focused on the assumption that a structural link is a necessary condition for an effective one, by studying the overlap between strong EC and SC links. In particular, our results indicate the importance of distinguishing between within and across network links, in relation to the degree of freedom given by the structural prior and its effect on model fit. A too strict structural prior negatively affected the model fit, since it does not allow the effective links to sufficiently diverge from it. While this is particularly relevant for the between network links and within high-order cortical areas, effective links within unimodal motor-sensory areas seem to align to the structural pathway independently of the prior restraint.

## Materials and Methods

### Data collection and preprocessing

A dataset of n=20 adult male C57BI6/J mice were previously acquired at the IIT laboratory (Italy). All in vivo experiments were conducted in accordance with the Italian law (DL 2006/2014, EU 63/2010, Ministero della Sanità, Roma) and the recommendations in the Guide for the Care and Use of Laboratory Animals of the National Institutes of Health. Animal research protocols were reviewed and consented by the animal care committee of the Italian Institute of Technology and Italian Ministry of Health. Animal preparation, image data acquisition and image data preprocessing for rsfMRI data have been described in greater detail elsewhere (Gutierrez-Barragan et al., 2019). Briefly, rsfMRI data were acquired on a 7.0-T scanner (Bruker BioSpin, Ettlingen) equipped with BGA-9 gradient set, using a 72-mm birdcage transmit coil, and a four-channel solenoid coil for signal reception. Single-shot BOLD echo planar imaging time series were acquired using an echo planar imaging sequence with the following parameters: repetition time/echo time, 1000/15 ms; flip angle, 30°; matrix, 100 ×100; field of view, 2.3 × 2.3 cm2; 18 coronal slices; slice thickness, 0.60 mm; 1920 volumes.

Image preprocessing has been previously described in greater detail elsewhere (Gutierrez-Barragan et al., 2022). Briefly, timeseries were despiked, motion corrected, skull stripped and spatially registered to an in-house EPI-based mouse brain template. Denoising and motion correction strategies involved the regression of mean ventricular signal plus 6 motion parameters (Rocchi et al., 2022). The resulting timeseries were band-pass filtered (0.01-0.1 Hz band) and then spatially smoothed with a Gaussian kernel of 0.5 mm full width at half maximum. After preprocessing, mean regional time-series were extracted for 74 (37+37) regions of interest (ROIs) derived from a predefined anatomical parcellation of the Allen Brain Institute (ABI), (Oh et al., 2014; Wang et al., 2020).

### Cortical network partitions

We partitioned the functional cortical networks into the lateral cortical network (LCN) (Liska et al., 2015), the default mode posterolateral network (DMNpost), the default mode midline network (DMNmid) and the salience (SAL) (Guitierrez-Barragan et al., 2022). In particular, LCN includes: primary and secondary motor, and primary and supplementary somatosensory areas. DMNpost contains: gustatory, posterior parietal association, temporal association and visceral areas. DMNmid contains: anterior circulate (dorsal and ventral), prelimbic, infralimbic, orbital and retrosplenial (agranular, ventral and dorsal) areas. SAL refers to the agranular insula areas (dorsal, posterior and ventral parts).

### Sparse DCM

The effective connectivity matrix was estimated at the single subject level using the method described in Prando et al. (2020), called sparse DCM. In line with the DCM framework, sparse DCM is a state-space model where the state is a system of differential equations representing the coupling among neural components, and the output model maps the neuronal activity to the measured BOLD signal through the haemodynamic response function (HRF). In particular, even if formulated in continuous time, sparse DCM operates in discrete time for the sake of computational efficiency. Moreover, to prevent spurious couplings, it includes a sparsity inducing prior in the EC estimation and a statistical linearization of the HRF model to deal with the high-dimensional solution space. Model inversion and parameter optimization are performed by an expectation-maximization (EM) algorithm.

A structurally-infomed variant of sparse DCM was used to evaluate the effect of imposing structural constraints on the EC matrix. Specifically, a subset of EC entries was forced to be zero in the inference process. Such a subset was selected from the SC matrix and it contained those links with a strength below a certain threshold. Three thresholds have been tested corresponding to the 60th, 40th and 20th percentiles of the SC entry distribution.

### Structural connectivity

The directed structural matrix was inferred from a directed weighted voxelwise model of the mouse brain (Knox et al., 2019) obtained through multiple viral microinjection experiments (Allen Mouse Brain Connectivity Atlas) and re-parceled into the same number of regions used for the rsfMRI data (Coletta et al., 2020).

### Structure-effective relationship

The structure-effective relationship was studied both at the global level and at the node level. In the first case, we considered how the strengths of all EC links distribute according to the structural connectome. In particular, we binarized the log-SC with different thresholds (from 10% to 90%, step 20) and used it to mask the EC. We refer to this as the global SC-EC coupling. Practically, we looked at the mean absolute EC strength of links both with and without a structural connection, MA(EC_strlink_) and MA(EC_nostrlink_) respectively, and normalized each mean with the global mean absolute EC, MA(EC). The same procedure was then repeated by reversing the role of the two measures, meaning thresholding the effective and evaluating the strength of the structure, i.e. global EC-SC coupling.

To study the relation at the node level, we estimated the coupling in each node for both incoming and outgoing links, i.e. by considering separately the rows and the columns of the connectivity matrices, respectively. In detail, for each node the coupling between structural and effective links was computed as the Spearman rank correlation between the top *k* entries of the effective and the related structural ones: we refer to this as the EC-SC coupling. Similarly, for each node the SC-EC coupling was computed as the Spearman rank between the top *k* entries of its structural vector and the corresponding effective entries. We will report results with *k*=15 that for our data corresponds to the 20% of ROIs (74) in the adopted parcelization (in Supplementary Materials, the coupling is shown under different thresholds, and statistical significance tested by random permutation test).

To quantify how much for a given node its strongest SC links deviate from its strongest EC links and vice-versa, we computed the *only SC* and *only EC* rates. The *only SC* rate is defined as the number of *only SC* over the total amount of considered links (*only SC* + *overlap*), where *only SC* is the number of structural edges which do not correspond to an effective edge and the *overlap* is the number of edges that are both structural and effective. Similarly for the *only EC* rate. A rate equals to 0 means that there are no lonely SC (or EC) edges; if it equals to 0.5 then lonely and overlapped edges are equally distributed; if it equals to 1 there is no overlap between effective and structural links. EC is a signed matrix with positive and negative entries modeling excitatory and inhibitory connections, respectively. In this work, we considered the strength of a connection independently from its sign, thus when we refer to EC we mean the absolute value of the actual DCM estimates.

### Goodness of fit metrics

After fitting a DCM model on each empirical single subject rsfMRI recordings, we used the model to generate 100 realizations of synthetic rsfMRI signals. The goodness of fit between empirical and simulated data was quantified by the correlation of their functional connectivity, i.e. the correlation between the triangular part of the empirical FC and simulated FC, and the similarity between the distributions of their dynamic FCs. In detail, to capture time-dependent properties of the data, the dynamic FC was computed using a sliding window of 50 s (with 25 s step), and the metric was given by the Kolmogorov-Smirnov distance between the triangular part of the empirical dFC and simulated dFC.

## Results

On each mouse rs-fMRI scan, we applied sparse DCM and obtained the related effective connectivity matrix. Fig. 1b shows the effective connectivity of a representative subject and, in panel a, the global directed structural matrix.

**Figure 1.**
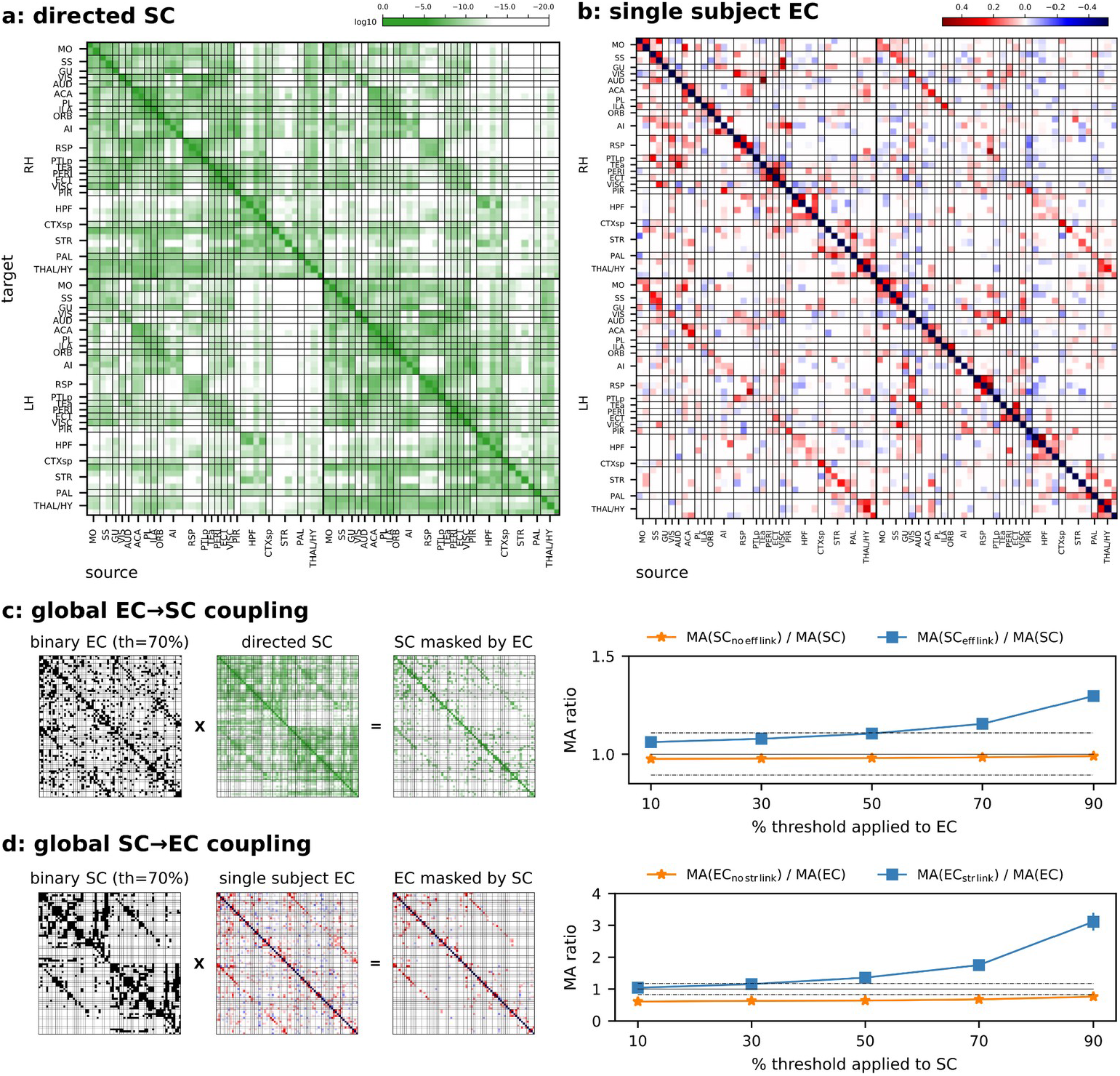
(a) The directed log-structural connectivity at the macro scale, obtained by a weighted voxel-wise model applied on the mouse axonal connectome. (b) Effective connectivity matrix of a representative subject, excitatory links are in red and inhibitory links in blues. In both matrices, target regions are on the y axis, and source regions on the x axis. Regions are grouped by hemisphere and sorted from cortical to subcortical areas. (c) EC->SC coupling: SC was masked by each single subject binary EC (given a certain threshold) and computed the related MA (mean absolute) ratios both for zero (orange line) and nonzero (blue line) mask entries. (d) SC->EC coupling, similarly to panel (c) with the mask derived from SC and applied on EC (for each subject and binarization thresholds). In both panels (c,d) an MA ratio greater than one on non-zero mask entries (blue lines) implies on average a high SC (EC) strength on nodes with high EC (SC) strength. As means of comparison, MA ratios computed within and across hemispheres are shown as dashed lines (>1 within, and <1 across).—

Firstly, we looked at the coupling between the two matrices from a global perspective. Thus we considered the mean absolute (MA) strength of the structural connectivity entries that are (and are not) associated with an effective link, under different binarization thresholds, and normalized with respect to the MA of the whole matrix, MA ratio (see Fig. 1c). We refer to this as EC-SC coupling: a binary EC mask was constructed to select the strongest links and then applied to the SC matrix. The same analysis was also repeated by reversing the role of the two measures, i.e. thresholding the structural and evaluating the strength of the effective entries (see Fig. 1d, SC-EC coupling). In both cases, the increase in the mean absolute ratio with increasing binarization threshold is evidence of a higher mean structural (effective) strength among nodes with the strongest effective (structural) links.

To characterize the relationship between the two types of connectivity at the node level, we quantified their coupling in each node, for both incoming and outgoing connections. The coupling between structural and effective links was computed as the Spearman rank correlation. Since sparse DCM was not structurally-informed, the coupling was evaluated when conditioned both on the effective (EC-SC coupling) and the structure (SC-EC coupling).

EC-SC coupling quantifies the relation between the strongest effective links and the related structural magnitude. In other words, it evaluates whether for a given node, its ordered effective vector is coherently supported by its structural vector. On the other hand, the SC-EC coupling quantifies how well the strongest structural links relate with the corresponding effective links in each node. In other words, if the effective strength follows the anatomical strength.

Due to the independence between SC and EC in our analysis, a match both in terms of link magnitude and ordering between SC and EC cannot be expected a priori. Thus, we firstly evaluated the EC-SC coupling to understand how the computed ECs resemble the SC in terms of agreement between the sorted effective links and the related structural ones for each node (see Fig. 2a). This is shown in Figure 2, where panels b and c contain the z-Spearman correlations across subjects with cortical nodes grouped by functional network for incoming and outgoing links, respectively, and in panel d at the single node-level. Importantly, when nodes were grouped by functional network and sorted by coupling strength, a unimodal-transmodal hierarchy emerged as previously described for the SC (Coletta et al., 2020). Specifically, EC links from and to transmodal networks, in particular SAL and DMN postero-lateral, were poorly coupled with the underlying structural links. On the contrary, in unimodal motor and somatosensory areas (LCN) coupling was higher. No clear-cut differences were found between incoming and outgoing links.

**Figure 2.**
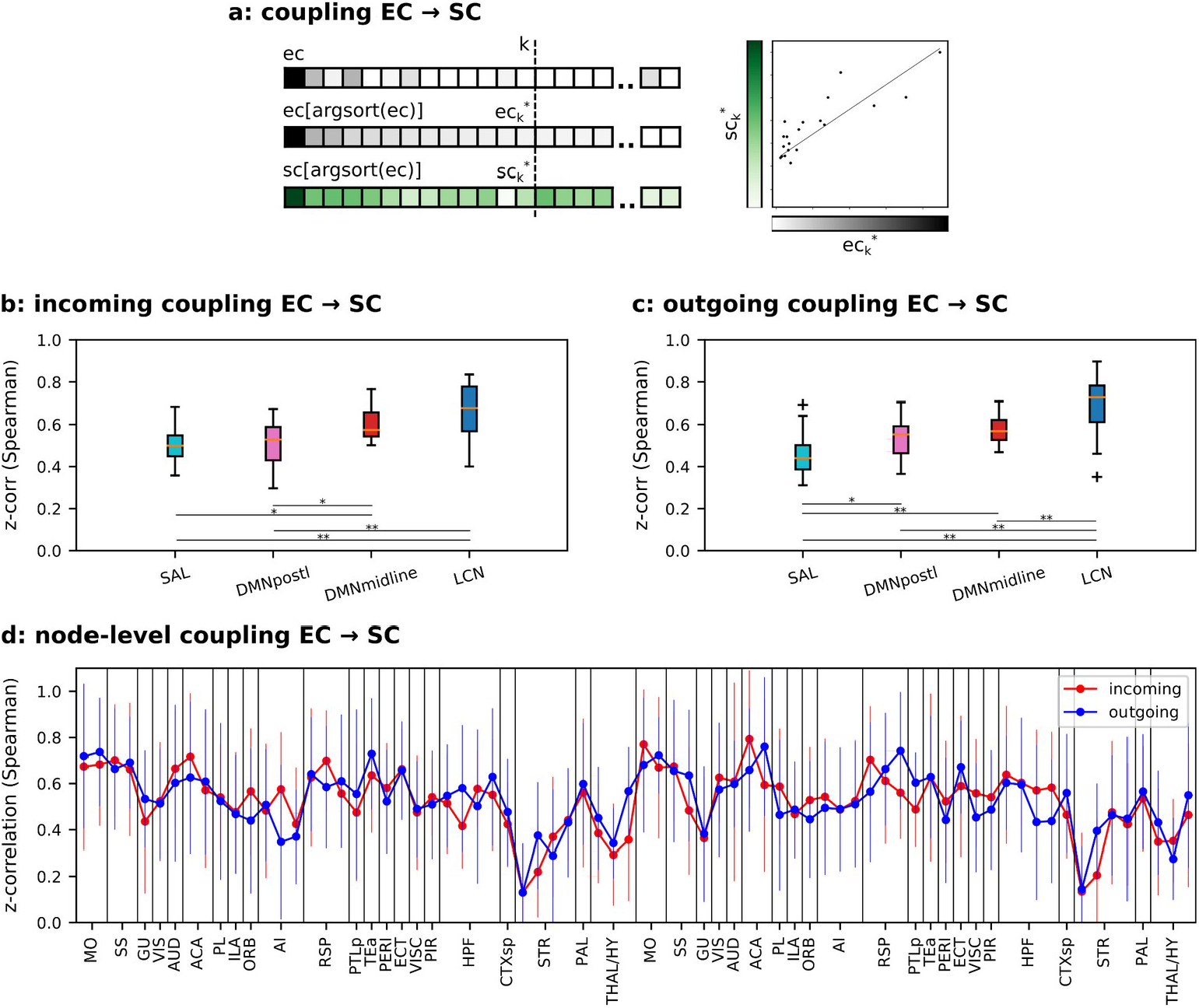
EC-SC coupling. (a) Explanatory cartoon of the EC-SC coupling computation for a given vector of links referred to the incoming or outgoing EC entries of a node (gray scale): the coupling is computed as the Spearman rank correlation between the strongest *k* EC entries and the related SC ones (green scale); argsort() gives the indices that would sort the vector. (b) Coupling across subjects computed on the incoming links and results grouped by functional networks with k=15. (c) Similar to panel (b) on outgoing links. *=p<0.05, **=p<0.01 and ***=p<0.001, ANOVA test with Tukey’s multiple comparison test. (d) EC-SC coupling at the node-level. No significant differences between incoming and outgoing node couplings have been detected. Paired t-test, Benjamini/Hochberg multiple testing correction.

Having reproduced a well-known functional hierarchy even with a no structurally informed algorithm reassured us on the validity of our model and prompted us to investigate SC-EC coupling.

To this aim, the strongest SC links were selected in each node, and correlated with the corresponding EC links. This is in line with the assumption that an effective link requires the presence of a structural link and it aims to test how the two measures are coupled.

Similarly to Fig. 2, Fig. 3 shows the coupling distribution across mice, with nodes grouped by their own functional network, separately for incoming and outgoing links, respectively, in panel b and c. Interestingly, we saw that SC-EC coupling did not follow the functional hierarchy as previously observed for EC-SC coupling. SAL network exhibited the lowest coupling, however, for DMN and LCN there was not a clear rank. Moreover, differently from the previous case, the coupling at the node-level (Fig. 3d) shows a significant decrease in the incoming links of the subcortical regions, i.e. striatum, pallidum and thalamus.

**Figure 3.**
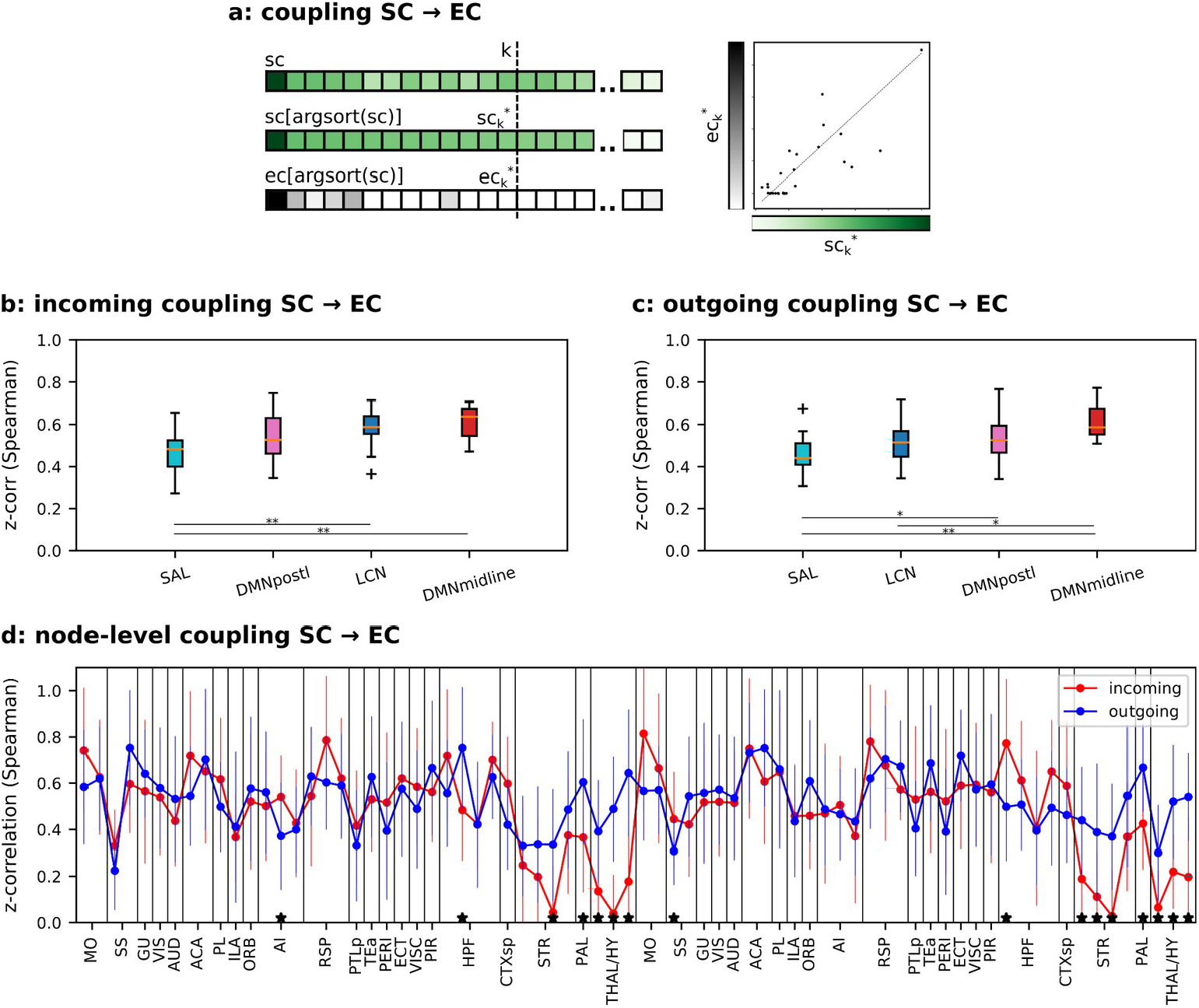
SC-EC coupling. (a) Explanatory cartoon of the SC-EC coupling computation for a given vector of links referred to the incoming or outgoing SC entries of a node (green scale): the coupling is computed as the Spearman rank correlation between the strongest *k* SC entries and the related EC ones (gray scale). (b) Coupling across subjects computed on the incoming links and results grouped by functional networks with k=15. (c) Similar to panel (b) on outgoing links. *=p<0.05, **=p<0.01 and ***=p<0.001, ANOVA test with Tukey’s multiple comparison test. (d) SC-EC coupling at the node-level. *=p<0.05, paired t-test, Benjamini/Hochberg multiple testing correction.

These inconsistencies between EC-SC and SC-EC couplings prompted further analysis to better understand how the set of strongest SC links differs from the set of strongest EC links. We firstly computed the overlap rate between the k-th strongest SC and EC links (note that this ratio tends to 1 with k getting closer to the network size, 74 in our data). Figure S3 shows the overlap ratio per network across different values of k (from 15, i.e. 20% of the nodes, to 55, i.e 74%, step 10). On average, with k=15 the ratio is 0.41 for the incoming links, and 0.42 for the outgoing links. The overlap ratio reaches average values around 0.75 with k=55.

This observation led us to focus on the strongest SC and EC links that do not overlap each other. In particular, the strongest SC links which do not correspond to a high effective connection, since they are at the origin of the inconsistencies between couplings.

For each cortical node, we firstly divided its strongest links (both SC and EC) into within-network and between-network links, and among them we counted the number of *only EC, only SC* and *overlap* links. The explanatory cartoon on the top of Fig. 4 focuses on within-network links, and it shows the example of a node with 2 *only SC* links, 1 *only EC* link and 3 *overlap* links. We used this information to compute the *only SC rate* as the ratio between the *only SC* and the sum of *overlap* and *only SC* links, Fig. 4a-b (and similarly the *only EC rate*, Fig. 4c-d). We found that the within-network *only SC rate* clearly changed between LCN and transmodal networks. In detail, LCN has a ratio significantly lower than 0.5, i.e. a small number of *only SC* links compared to the *overlap* links, while transmodal networks have an *only SC* rate closer to 0.5, i.e. a comparable number of *only SC* links and *overlap* ones. From the side of the strongest EC links, the *only EC rate* is always smaller than 0.5 meaning that the majority of effective links are supported by a structural link (Fig. 4c-d). Transparent bars refer to the structurally-informed DCM results with thresholds of 60%, 40% and 20%, respectively (percentage of strongest kept links). As expected, the rate tends to 0 with the percentage of kept links getting lower.

**Figure 4.**
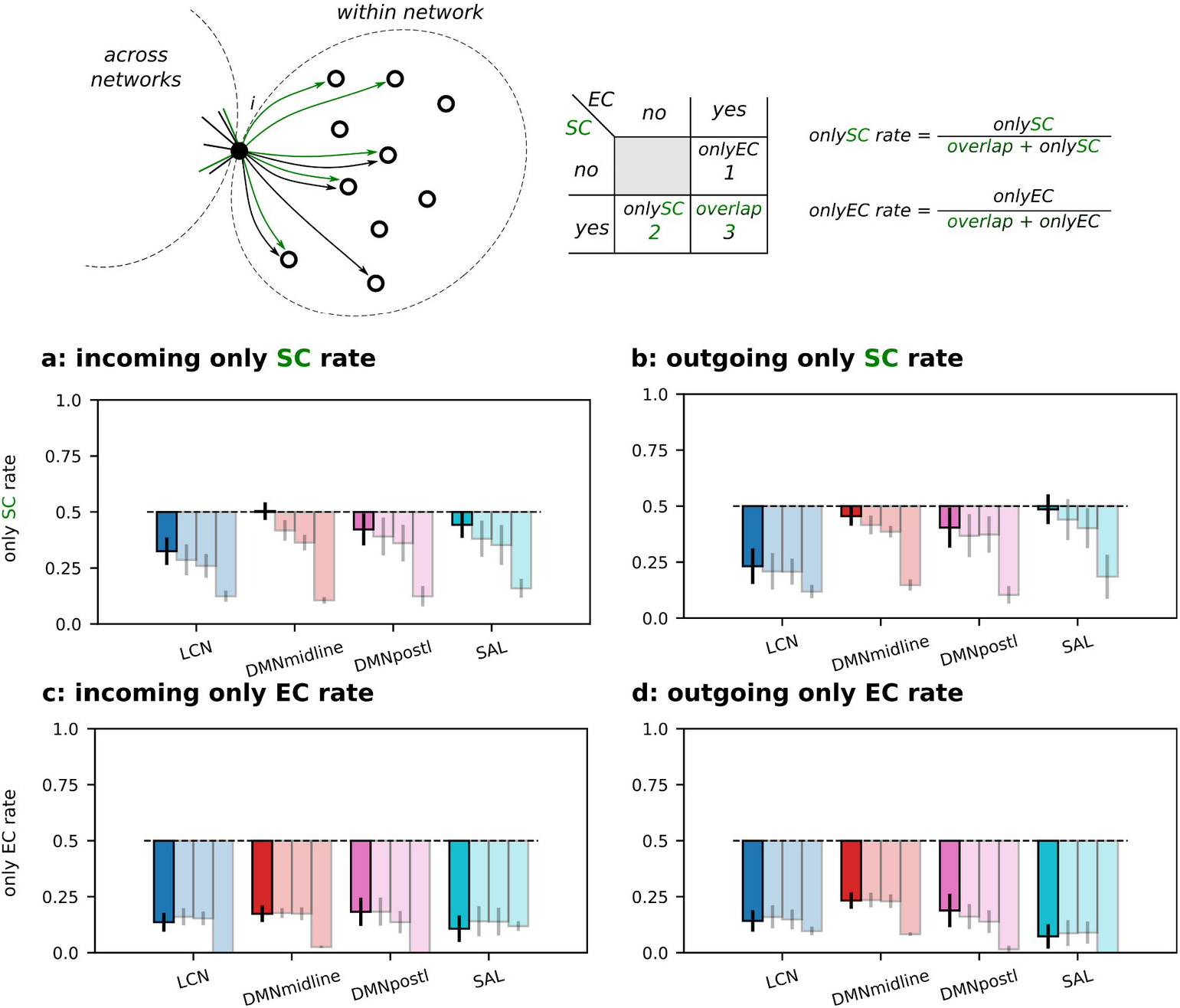
*Only SC* and *only EC* rates within networks. Explanatory cartoon of the *only SC* and *EC* rates computed on the within-network outgoing links of the given node *i*, SC links are in green and EC links in black. (a) Incoming *only SC rate*. (b) Outgoing *only SC rate*. (c) Incoming *only EC rate*. (d) Outgoing *only EC rate*. All ratios were computed at the node level and results grouped by functional network. Transparent bars refer to the structurally-informed DCM results with thresholds of 60%, 40% and 20%, respectively (percentage of kept links).

We repeated the analysis by looking at across network links. Both *only SC* and *EC rates* are greater than 0.5 by using non-structurally informed DCM (and for both incoming and outgoing links, see Fig. 5a-d full-color bars). This means a low overlap between strong EC and SC links. When we limited DCM to the strongest SC links, the overlap increased with the decrease in the percentage of kept links, see Fig. 5a-d transparent color bars. In particular, *only SC/EC rates* dropped below 0.5 when EC was limited to the 20% strongest SC links.

**Figure 5.**
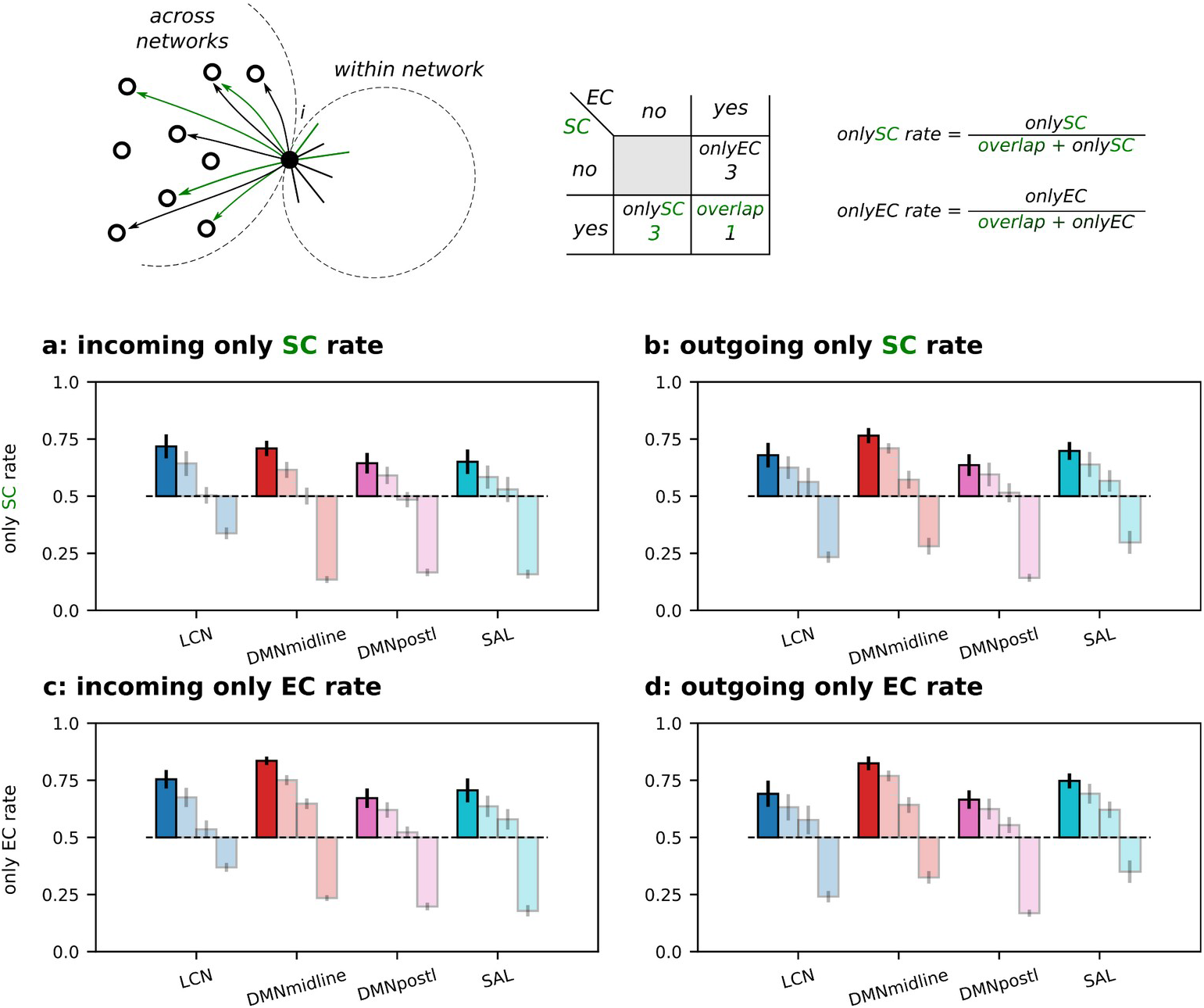
*Only SC* and *only EC* rates across networks. Explanatory cartoon of the *only SC* and *EC* rates computed on the across network outgoing links of the given node *i*, SC links are in green and EC links in black. (a) Incoming *only SC rate*. (b) Outgoing *only SC rate*. (c) Incoming *only EC rate*. (d) Outgoing *only EC rate*. All ratios were computed at the node level and results grouped by functional network. Transparent bars refer to the structurally-informed DCM results with thresholds of 60%, 40% and 20%, respectively (percentage of kept links).

The impact of a structural prior on the goodness of fit to the empirical data is shown in Fig. 6. We computed the variation of log-likelihood for each subject with the non-structurally informed DCM as reference (Fig. 6a). On average, it shows a reduction of the log-likelihood if a structural constraint was added. Moreover, we used DCM to generate 100 realizations of BOLD data for each subject and prior condition. We computed the correlation between the empirical and simulated FC matrices (Fig. 6b, left y-axis), and the Kolmogorov-Smirnov distance between dynamic FCs (Fig. 6b, right y-axis). Altogether, these results show that the most severe structural constraint (20% SC) had the highest impact on the goodness of fit causing the strongest reduction in *only SC/EC rates* (across networks this is the only case where rates became less than 0.5).

**Figure 6.**
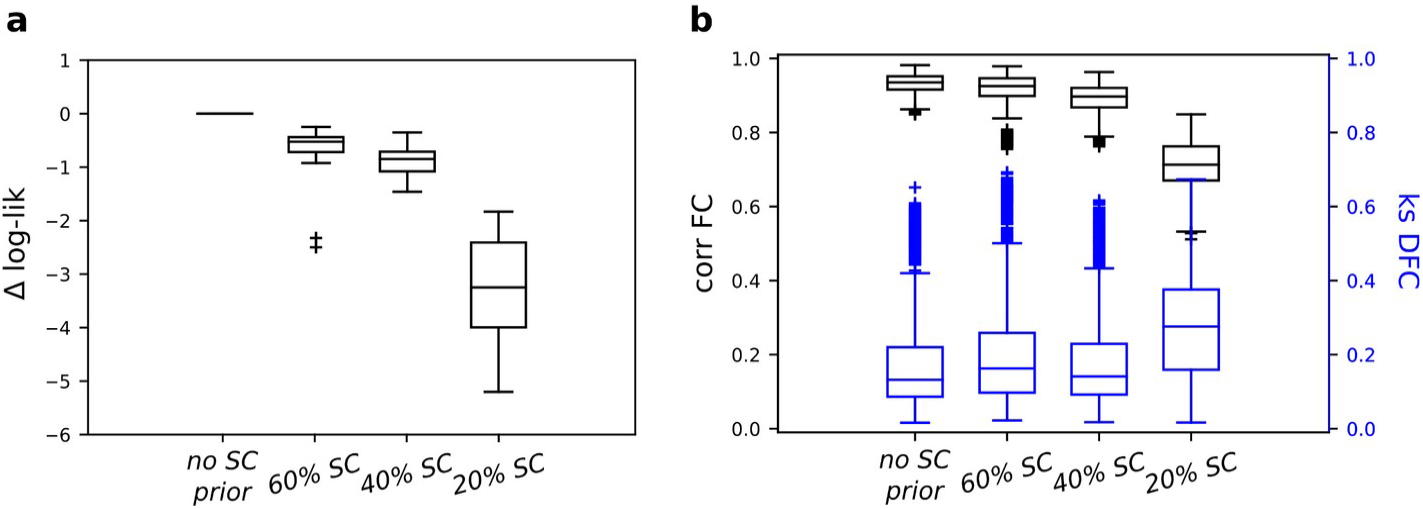
Model fit quantified as: (a) the log-likelihood difference between the structurally informed DCMs and the non-structurally informed DCM (used as reference, at the subject level); (b) the capability of generating data similar to the empirical recordings used to fit the DCM model, in terms of correlation between their FCs (left y-axis) and KS distance of their dynamic FCs (right y-axis), for non-structurally informed DCM and structurally informed DCM with different SC thresholds, i.e. 60, 40 and 20 (percentage of kept links).

Adding a structural prior also affects both couplings computed above, i.e. EC-SC and SC-EC couplings. Structurally informed DCM gave the expected functional rank in both couplings. The higher the constraint imposed by the prior, the more the functional hierarchy became unmistakable across functional networks, see Fig. S2. Interestingly, this is true even for the SC-EC coupling that did not give the expected unimodal-transmodal hierarchy in the non-structurally informed case, Fig. 3b-c. Moreover, the rank is more pronounced with the incoming coupling in both EC-SC and SC-EC (Fig. S2a/c).

## Discussion

The relation between structural and functional properties of brain connectivity is a central topic in neuroscience. Previous studies have shown the lack of a one-to-one correspondence between structural and functional connectivity. In line with this, a branch of literature has rapidly grown on local structure-functional coupling showing a strong dependence with cortical hierarchies (Vázquez-Rodríguez et al., 2019; Preti and Ville, 2019; Liu et al., 2022b). Most of these results were obtained on human data by studying the relation between FC and SC. Here, we employed the mouse brain model for which both an accurate reconstruction of the axonal paths (Coletta et al., 2020; Harris et al., 2019), and a consolidated protocol to acquire resting-state fMRI data are available (Liska et al., 2015). Moreover, the recent developments in the framework of dynamical causal models (DCM) have given variants of DCM suited to compute effective connectivity (EC) in whole-brain networks (Razi et al., 2017; Frässle et al., 2021; Prando et al., 2020). In the context of DCM, the advantage given by structurally-informed prior has been largely proved by tuning the prior variance of each effective link proportionally to the likelihood that such link anatomically exists (Stephan et al., 2009; Sokolov et al., 2019).

Here, we started with a non-structurally informed DCM and compared each single mouse EC with the mouse structural connectome obtained at the population level. We found that a higher mean structural strength corresponds to a higher mean effective strength, and vice versa. This is consistent with prior studies showing how DCM benefits from structural information (Crimi et al., 2021). However, our aim was to further our understanding of coupling between EC and SC at the node level, and characterize their overlap.

Importantly we found that EC-SC coupling, i.e. the coupling driven by the strongest effective links, follows the unimodal-transmodal functional hierarchy at the cortical level previously identified in the mouse (Coletta et al., 2020). This relation replicates and expands human findings, by showing that the relationship previously observed in humans is largely driven by EC-SC, and not vice versa (Baum et al., 2020; Esfahlani et al., 2021; Gu et al., 2021). In this respect, the coupling of the salience (SAL) network is of particular interest, as both the incoming and outgoing connections of the SAL have the lowest coupling strength compared to the other functional networks. This is in line with what was previously reported by Liu et al. (2022b), who found that SAL is one of the most dynamic networks in terms of structure-functional coupling.

Interestingly, SC-EC coupling (i.e. the coupling driven by the strongest structural links) did not replicate the same cortical hierarchy found with EC-SC, meaning that when conditioning on the structure, links that do not meet the expected relationship were included. This prompted us to focus on the overlap between EC and SC. Our results show two different scenarios, depending on whether we consider within or between network links. In particular, within high-order cortical areas we found strong structural connections that did not exhibit a correspondingly strong effective link. The mismatch between SC and EC was even more clear on links across networks in which also the majority of strong effective links were not supported by a structural one. Thus, only the connections within unimodal sensory motor networks aligned both in terms of effective and structural strength, highlighting the different degree of segregation of each network. This finding is consistent with the fact that the unimodal somatomotor network is densely connected and spatially close, while this is not the case for the DMN or higher transmodal networks (Suárez et al., 2020; Whitesell et al., 2021)

A different perspective to look at the heterogeneity of structural and functional connections considers how communication occurs in the network (Pedersen et al., 2020). The specific link organization within and between networks has been recently explained by considering the preferred pattern of structural connections that each network adopts to communicate (Bazinet et al., 2021; Liu et al., 2022a). By rephrasing our results in these terms, the stronger structure-functional coupling in unimodal regions is a consequence of the preferred local scale, i.e. monosynaptic links, on which communication occurs. Moving along the hierarchy toward the transmodal cortex, the optimal scale globally extends involving more polysynaptic pathways. Therefore, our observations of a different connection profile across unimodal and polymodal networks in terms of SC and EC might be explained with a gradient of optimal scales ranging from local (within LCN, where SC and EC overlap), to a more global scale occurring between networks.

Additional evidence on the effect of monosynaptic vs. polysynaptic paths is that the SC-EC coupling significantly decreased in the incoming links of some subcortical nodes, i.e. lateral septal complex, pallidum, thalamus and hypothalamus. This finding is consistent with previous observation of a higher mismatch between FC and SC in subcortical networks due to the polysynaptic nature of their anatomical links (Grandjean et al., 2017).

In an attempt to better understand the coupling between SC and EC, we also included a structural constraint on DCM. Specifically, we forced EC to be always supported by a structural link, so we repeated the inference for different percentiles of kept structural connections. Following the previous interpretation of multiscale communication, by constraining EC to SC, we limited the spectrum of the spatial scale to be more local as the percentage of kept links decreases. This analysis yielded three results: firstly, as expected the overlap between strong EC and SC links was found to be increased, i.e. the *only SC* and *EC* ratios as shown in Fig.s 4 and 5, tended to zero both within and across networks. Secondly, the previously observed functional hierarchy became more evident when EC was forced to comply with SC (Supplementary Fig. S2) both when computed through EC-SC coupling and SC-EC coupling. In particular, incoming effective links, when strongly constrained by the structure, revealed a well distinct cortical rank that reflects the expected hierarchy. This was mainly due to LCN nodes in which their effective links strongly coupled with their structural ones. The same effect was not observed in outgoing links. This finding is in line with the results of Sokolov et al. (2020), who showed that DCM with a prior built on incoming structural information outperformed models informed by outgoing structural information as well as those without structural information. Lastly, the third finding relates to the fact that DCM is a generative model so it can produce new realizations of the same dynamical system. Within this framework, we tested how the capability of our model to generate data as similar as possible to the empirical ones, varied in relation to a structural constraint. Our results showed a reduction of the goodness of fit (both in terms of fit to the empirical signals and capability to reproduce the empirical static and dynamic functional connectivity) when the structural constraint was too strict. Here, it is important to note how the overlap between SC and EC got higher when the structural prior became stricter in DCM, in line with the result of Bazinet et al. (2021). Altogether these results suggest that a structural constraint on DCM should consider the heterogeneity of the EC/SC coupling: it can be more severe on links within sensory motor network, whereas a higher degree of freedom is needed especially on links across networks.

The interpretation of these results must also take into account some limitations. First, while using the murine model, we leveraged the possibility of differentiating incoming and outgoing SC connections. However, it is unclear how well these results apply to higher mammalian species characterized by larger white matter tracts, and by proportionally denser long-range connectivity (Mota et al., 2019). Second, by using the mouse brain connectome, i.e. a single structural matrix, the inter-subject variability could not be considered, while from recent studies interindividual variability in human (Gu et al., 2021), along with other factors, e.g. time dependence (Liu et al., 2022b), and receptor maps (Hansen et al., 2022), have shown to play a role in relating structure to function.

In summary, the present work studies the relation between SC and EC in terms of their coupling and overlap by conditioning both on the strongest SC and EC links. The novelty of our work concerns mainly three aspects: i) it brings EC into the field of structural-functional coupling, ii) it differentiates between EC-SC and SC-EC couplings (by conditioning it on the strongest EC and SC links, respectively) highlighting how non-structurally informed EC and SC differ, and iii) it characterizes the role of a structurally informed prior in DCM in terms of both its effect on the relation between SC and EC and the generative capability of the model. Briefly, we found that when conditioning on the strongest EC links, the coupling follows the unimodal-transmodal functional hierarchy. Whereas the reverse is not true, indeed there are strong SC links within high-order cortical areas which do not correspond to a strong EC link. This mismatch is even more clear across networks. Only the connections within sensory motor networks freely align both in terms of effective and structural strength.

## Acknowledgements

This research was supported by the DEI Proactive grant “Personalized whole brain models for neuroscience: inference and validation” from the Department of Information Engineering of the University of Padova (Italy). A. Gozzi is supported by the European Research Council (ERC-DISCONN, No. 802371), NIH (1R21MH116473-01A1) and the Telethon foundation (GGP19177).

## Supplementary Materials

**Figure S1.**
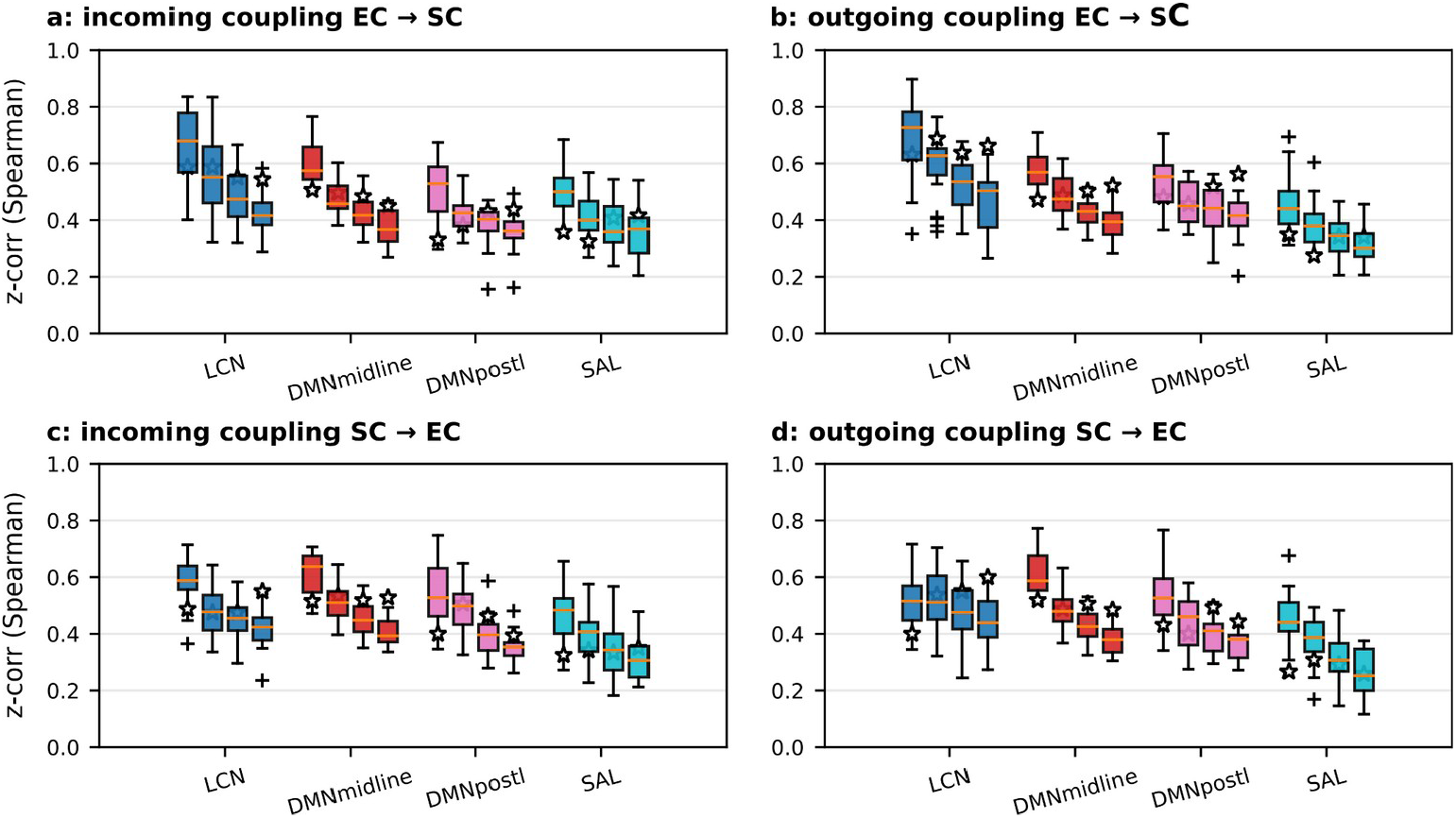
(a-b) EC-SC coupling, an expanded version of Figure 2b-c across different values of threshold k (k=15, 20, 25, 30) that is the number of top EC entries selected in each node incoming and outgoing links, on which the Spearman rank correlation was computed with the corresponding SC entries. (c-d) SC-EC coupling, similar to Figure 3b-c, extended by increasing the threshold k from 15 to 30 (step 5), as before. The first boxplot in each quartet refers to k=15, thus equals to what reported in the main text. In each boxplot, white start marker shows the average rate of nodes with significant correlation in that functional network. Significance was computed by random permutations test (p=0.05, n=500). Overall, it shows a decrease in the coupling when k increases, and also a higher EC-SC coupling in LCN which makes the gradient from unimodal to transmodal networks more evident.

**Figure S2.**
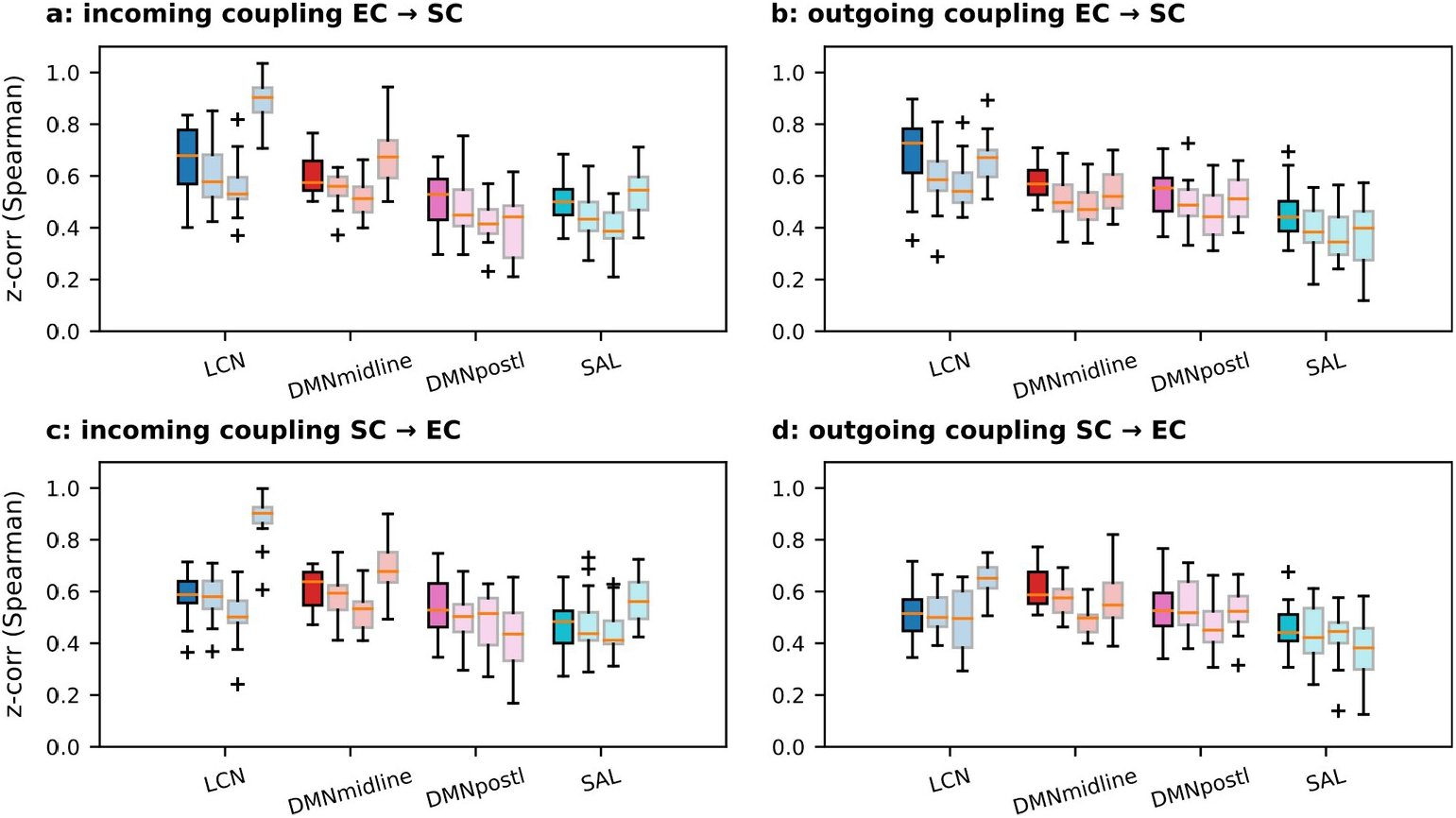
(a-b) EC-SC coupling, similar to Figure 2b-c, extended with the couplings computed on the results from the structurally-informed DCM with different SC thresholds, i.e. 60, 40 and 20% (percentage of kept links). (c-d) SC-EC coupling, similar to Figure 3b-c, extended with the structurally informed results, as before.

**Figure S3.**
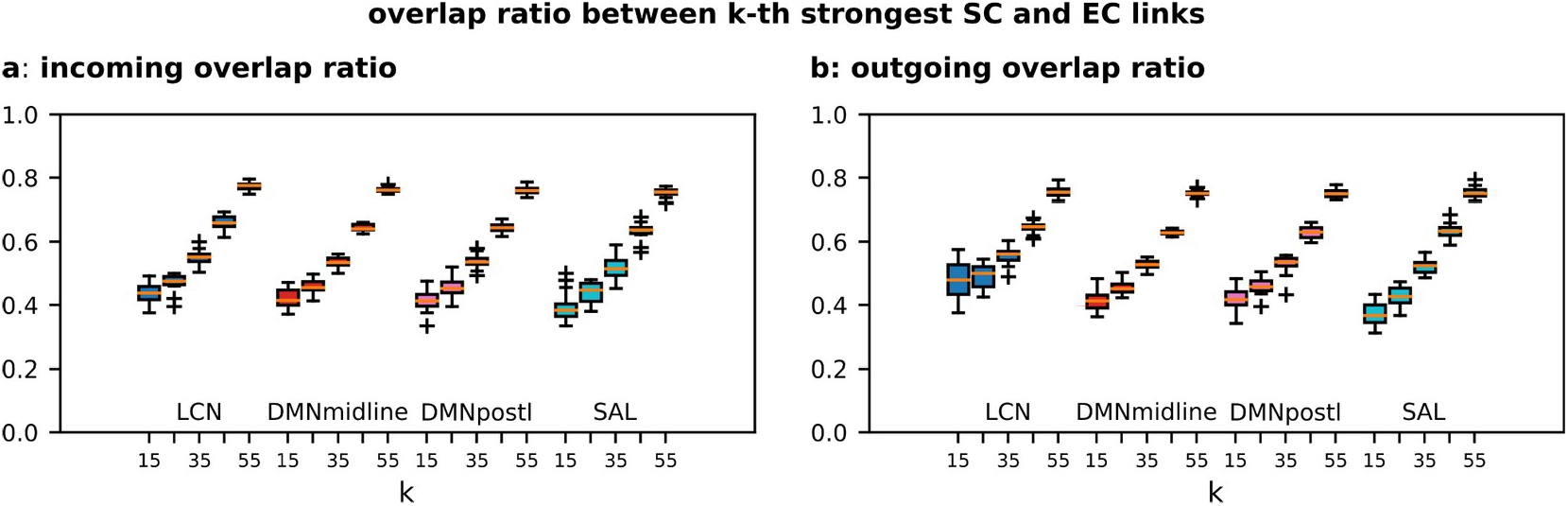
Overlap ratio between k-th strongest SC and EC links computed on each node and grouped by functional networks, k from 15 to 55 (step 10).

## References

Atasoy, S., Donnelly, I., Pearson, J., 2016. Human brain networks function in connectome-specific harmonic waves. Nature Communications 2016 7:1 7, 1–10. doi:10.1038/ncomms10340.

Avena-Koenigsberger, A., Misic, B., Sporns, O., 2017. Communication dynamics in complex brain networks. Nature reviews. Neuroscience 19, 17–33. doi:10.1038/NRN.2017.149.

Baum, G.L., Cui, Z., Roalf, D.R., Ciric, R., Betzel, R.F., Larsen, B., Cieslak, M., Cook, P.A., Xia, C.H., Moore, T.M., Ruparel, K., Oathes, D.J., Alexander-Bloch, A.F., Shinohara, R.T., Raznahan, A., Gur, R.E., Gur, R.C., Bassett, D.S., Satterthwaite, T.D., 2020. Development of structure–function coupling in human brain networks during youth. Proceedings of the National Academy of Sciences of the United States of America 117, 771–778. doi:10.1073/PNAS.1912034117/SUPPL_FILE/PNAS.1912034117.SAPP.PDF.

Bazinet, V., de Wael, R.V., Hagmann, P., Bernhardt, B.C., Misic, B., 2021. Multiscale communication in cortico-cortical networks. NeuroImage 243. doi:10.1016/J.NEUROIMAGE.2021.118546.

Bullmore, E., Sporns, O., 2009. Complex brain networks: graph theoretical analysis of structural and functional systems. Nature Reviews Neuroscience 2009 10:3 10, 186–198. doi:10.1038/nrn2575.

Chen, Y., Bukhari, Q., Lin, T.W., Sejnowski, T.J., 2022. Functional connectivity of fmri using differential covariance predicts structural connectivity and behavioral reaction times. Network Neuroscience 6, 614–633. doi:10.1162/NETN_A_00239.

Coletta, L., Pagani, M., Whitesell, J.D., Harris, J.A., Bernhardt, B., Gozzi, A., 2020. Network structure of the mouse brain connectome with voxel resolution. Science advances 6. doi:10.1126/SCIADV.ABB7187.

Crimi, A., Dodero, L., Sambataro, F., Murino, V., Sona, D., 2021. Structurally constrained effective brain connectivity. NeuroImage 239, 118288. doi:10.1016/J.NEUROIMAGE.2021.118288.

D’Angelo, E., Jirsa, V., 2022. The quest for multiscale brain modeling. Trends in neurosciences 45, 777–790. doi:10.1016/J.TINS.2022.06.007.

Esfahlani, F.Z., Bertolero, M.A., Bassett, D.S., Betzel, R.F., 2020. Space-independent community and hub structure of functional brain networks. NeuroImage 211, 116612. doi:10.1016/J.NEUROIMAGE.2020.116612.

Friston, K.J., Harrison, L., Penny, W., 2003. Dynamic causal modelling. Neuroimage 19, 1273–1302. doi:10.1016/S1053-8119(03)00202-7.

Frässle, S., Harrison, S.J., Heinzle, J., Clementz, B.A., Tamminga, C.A., Sweeney, J.A., Gershon, E.S., Keshavan, M.S., Pearlson, G.D., Powers, A., Stephan, K.E., 2021. Regression dynamic causal modeling for resting-state fmri. Human brain mapping 42, 2159–2180. doi:10.1002/HBM.25357.

Grandjean, J., Zerbi, V., Balsters, J.H., Wenderoth, N., Rudin, M., 2017. Structural basis of large-scale functional connectivity in the mouse. The Journal of neuroscience: the official journal of the Society for Neuroscience 37, 8092–8101. doi:10.1523/JNEUROSCI.0438-17.2017.

Gu, S., Fotiadis, P., Parkes, L., Xia, C.H., Gur, R.C., Gur, R.E., Roalf, D.R., Satterthwaite, T.D., Bassett, D.S., 2022. Network controllability mediates the relationship between rigid structure and flexible dynamics. Network Neuroscience 6, 275–297. doi:10.1162/NETN_A_00225.

Gu, S., Pasqualetti, F., Cieslak, M., Telesford, Q.K., Yu, A.B., Kahn, A.E., Medaglia, J.D., Vettel, J.M., Miller, M.B., Grafton, S.T., Bassett, D.S., 2015. Controllability of structural brain networks. Nature Communications 2015 6:1 6, 1–10. doi:10.1038/ncomms9414.

Gu, Z., Jamison, K.W., Sabuncu, M.R., Kuceyeski, A., 2021. Heritability and interindividual variability of regional structure-function coupling. Nature Communications 2021 12:1 12, 1–12. doi:10.1038/s41467-021-25184-4.

Gutierrez-Barragan, D., Basson, M.A., Panzeri, S., Gozzi, A., 2019. Infraslow state fluctuations govern spontaneous fmri network dynamics. Current Biology 29. doi:10.1016/j.cub.2019.06.017.

Gutierrez-Barragan, D., Singh, N.A., Alvino, F.G., Coletta, L., Rocchi, F., Guzman, E.D., Galbusera, A., Uboldi, M., Panzeri, S., Gozzi, A., 2022. Unique spatiotemporal fmri dynamics in the awake mouse brain. Current biology: CB 32, 631–644.e6. doi:10.1016/J.CUB.2021.12.015.

Hansen, J.Y., Shafiei, G., Markello, R.D., Smart, K., Cox, S.M., Nørgaard, M., Beliveau, V., Wu, Y., Gallezot, J.D., Étienne Aumont, Servaes, S., Scala, S.G., DuBois, J.M., Wainstein, G., Bezgin, G., Funck, T., Schmitz, T.W., Spreng, R.N., Galovic, M., Koepp, M.J., Duncan, J.S., Coles, J.P., Fryer, T.D., Aigbirhio, F.I., McGinnity, C.J., Hammers, A., Soucy, J.P., Baillet, S., Guimond, S., Hietala, J., Bedard, M.A., Leyton, M., Kobayashi, E., Rosa-Neto, P., Ganz, M., Knudsen, G.M., Palomero-Gallagher, N., Shine, J.M., Carson, R.E., Tuominen, L., Dagher, A., Misic, B., 2022. Mapping neurotransmitter systems to the structural and functional organization of the human neocortex. Nature Neuroscience 2022 25:11 25, 1569–1581. doi:10.1038/s41593-022-01186-3.

Harris, J.A., Mihalas, S., Hirokawa, K.E., Whitesell, J.D., Choi, H., Bernard, A., Bohn, P., Caldejon, S., Casal, L., Cho, A., Feiner, A., Feng, D., Gaudreault, N., Gerfen, C.R., Graddis, N., Groblewski, P.A., Henry, A.M., Ho, A., Howard, R., Knox, J.E., Kuan, L., Kuang, X., Lecoq, J., Lesnar, P., Li, Y., Luviano, J., McConoughey, S., Mortrud, M.T., Naeemi, M., Ng, L., Oh, S.W., Ouellette, B., Shen, E., Sorensen, S.A., Wakeman, W., Wang, Q., Wang, Y., Williford, A., Phillips, J.W., Jones, A.R., Koch, C., Zeng, H., 2019. Hierarchical organization of cortical and thalamic connectivity. Nature 2019 575:7781 575, 195–202. doi:10.1038/s41586-019-1716-z.

Honey, C.J., Thivierge, J.P., Sporns, O., 2010. Can structure predict function in the human brain? NeuroImage 52, 766–776. doi:10.1016/J.NEUROIMAGE.2010.01.071.

Kale, P., Zalesky, A., Gollo, L.L., 2018. Estimating the impact of structural directionality: How reliable are undirected connectomes? Network Neuroscience 2, 259. doi:10.1162/NETN_A_00040.

Knox, J.E., Harris, K.D., Graddis, N., Whitesell, J.D., Zeng, H., Harris, J.A., Shea-Brown, E., Mihalas, S., 2019. High-resolution data-driven model of the mouse connectome. Network Neuroscience 3. doi:10.1162/netn_a_00066.

Li, G., Yap, P.T., 2022. From descriptive connectome to mechanistic connectome: Generative modeling in functional magnetic resonance imaging analysis. Frontiers in Human Neuroscience 16, 578. doi:10.3389/FNHUM.2022.940842/BIBTEX.

Liska, A., Galbusera, A., Schwarz, A.J., Gozzi, A., 2015. Functional connectivity hubs of the mouse brain. NeuroImage 115, 281–291. doi:10.1016/J.NEUROIMAGE.2015.04.033.

Liu, Z.Q., Betzel, R.F., Misic, B., 2022a. Benchmarking functional connectivity by the structure and geometry of the human brain. Network Neuroscience 6, 937–949. doi:10.1162/NETN_A_00236.

Liu, Z.Q., Vázquez-Rodríguez, B., Spreng, R.N., Bernhardt, B.C., Betzel, R.F., Misic, B., 2022b. Time-resolved structurefunction coupling in brain networks. Communications Biology 2022 5:1 5, 1–10. doi:10.1038/s42003-022-03466-x.

Lynn, C.W., Bassett, D.S., 2019. The physics of brain network structure, function and control. Nature Reviews Physics 2019 1:5 1, 318–332. doi:10.1038/s42254-019-0040-8.

Maier-Hein, K.H., Neher, P.F., Houde, J.C., Côté, M.A., Garyfallidis, E., Zhong, J., Chamberland, M., Yeh, F.C., Lin, Y.C., Ji, Q., Reddick, W.E., Glass, J.O., Chen, D.Q., Feng, Y., Gao, C., Wu, Y., Ma, J., Renjie, H., Li, Q., Westin, C.F., Deslauriers-Gauthier, S., González, J.O.O., Paquette, M., St-Jean, S., Girard, G., Rheault, F., Sidhu, J., Tax, C.M., Guo, F., Mesri, H.Y., Dávid, S., Froeling, M., Heemskerk, A.M., Leemans, A., Boré, A., Pinsard, B., Bedetti, C., Desrosiers, M., Brambati, S., Doyon, J., Sarica, A., Vasta, R., Cerasa, A., Quattrone, A., Yeatman, J., Khan, A.R., Hodges, W., Alexander, S., Romascano, D., Barakovic, M., Auría, A., Esteban, O., Lemkaddem, A., Thiran, J.P., Cetingul, H.E., Odry, B.L., Mailhe, B., Nadar, M.S., Pizzagalli, F., Prasad, G., Villalon-Reina, J.E., Galvis, J., Thompson, P.M., Requejo, F.D.S., Laguna, P.L., Lacerda, L.M., Barrett, R., Dell’Acqua, F., Catani, M., Petit, L., Caruyer, E., Daducci, A., Dyrby, T.B., Holland-Letz, T., Hilgetag, C.C., Stieltjes, B., Descoteaux, M., 2017. The challenge of mapping the human connectome based on diffusion tractography. Nature Communications 2017 8:1 8, 1–13. doi:10.1038/s41467-017-01285-x.

Margulies, D.S., Ghosh, S.S., Goulas, A., Falkiewicz, M., Huntenburg, J.M., Langs, G., Bezgin, G., Eickhoff, S.B., Castellanos, F.X., Petrides, M., Jefferies, E., Smallwood, J., 2016. Situating the default-mode network along a principal gradient of macroscale cortical organization. Proceedings of the National Academy of Sciences of the United States of America 113, 12574–12579. doi:10.1073/PNAS.1608282113/SUPPL_FILE/PNAS.201608282SI.PDF.

Miŝic, B., Betzel, R.F., Reus, M.A.D., Heuvel, M.P.V.D., Berman, M.G., McIntosh, A.R., Sporns, O., 2016. Network-level structure-function relationships in human neocortex. Cerebral Cortex 26, 3285–3296. doi:10.1093/CERCOR/BHW089.

Mota, B., Santos, S.E.D., Ventura-Antunes, L., Jardim-Messeder, D., Neves, K., Kazu, R.S., Noctor, S., Lambert, K., Bertelsen, M.F., Manger, P.R., Sherwood, C.C., Kaas, J.H., Herculano-Houzel, S., 2019. White matter volume and white/gray matter ratio in mammalian species as a consequence of the universal scaling of cortical folding. Proceedings of the National Academy of Sciences of the United States of America 116, 15253–15261. doi:10.1073/PNAS.1716956116/SUPPL_FILE/PNAS.1716956116.SD01.DOCX.

Oh, S.W., Harris, J.A., Ng, L., Winslow, B., Cain, N., Mihalas, S., Wang, Q., Lau, C., Kuan, L., Henry, A.M., Mortrud, M.T., Ouellette, B., Nguyen, T.N., Sorensen, S.A., Slaughterbeck, C.R., Wakeman, W., Li, Y., Feng, D., Ho, A., Nicholas, E., Hirokawa, K.E., Bohn, P., Joines, K.M., Peng, H., Hawrylycz, M.J., Phillips, J.W., Hohmann, J.G., Wohnoutka, P., Gerfen, C.R., Koch, C., Bernard, A., Dang, C., Jones, A.R., Zeng, H., 2014. A mesoscale connectome of the mouse brain. Nature 2014 508:7495 508, 207–214. doi:10.1038/nature13186.

Pedersen, M., Omidvarnia, A., Shine, J.M., Jackson, G.D., Zalesky, A., 2020. Reducing the influence of intramodular connectivity in participation coefficient. Network Neuroscience 4, 416–431. doi:10.1162/NETN_A_00127.

Prando, G., Zorzi, M., Bertoldo, A., Corbetta, M., Zorzi, M., Chiuso, A., 2020. Sparse dcm for whole-brain effective connectivity from resting-state fmri data. NeuroImage 208. doi:10.1016/j.neuroimage.2019.116367.

Preti, M.G., Ville, D.V.D., 2019. Decoupling of brain function from structure reveals regional behavioral specialization in humans. Nature Communications 2019 10:1 10, 1–7. doi:10.1038/s41467-019-12765-7.

Razi, A., Seghier, M.L., Zhou, Y., McColgan, P., Zeidman, P., Park, H.J., Sporns, O., Rees, G., Friston, K.J., 2017. Large-scale dcms for resting-state fmri. Network neuroscience (Cambridge, Mass.) 1, 222–241. doi:10.1162/NETN_A_00015.

Ritter, P., Schirner, M., Mcintosh, A.R., Jirsa, V.K., 2013. The virtual brain integrates computational modeling and multimodal neuroimaging. Brain connectivity 3, 121–145. doi:10.1089/BRAIN.2012.0120.

Rocchi, F., Canella, C., Noei, S., Gutierrez-Barragan, D., Coletta, L., Galbusera, A., Stuefer, A., Vassanelli, S., Pasqualetti, M., Iurilli, G., Panzeri, S., Gozzi, A., 2022. Increased fmri connectivity upon chemogenetic inhibition of the mouse prefrontal cortex. Nature Communications 2022 13:1 13, 1–15. doi:10.1038/s41467-022-28591-3.

Schirner, M., Kong, X., Yeo, B.T., Deco, G., Ritter, P., 2022. Dynamic primitives of brain network interaction. NeuroImage 250, 118928. doi:10.1016/J.NEUROIMAGE.2022.118928.

Sokolov, A.A., Zeidman, P., Erb, M., Ryvlin, P., Pavlova, M.A., Friston, K.J., 2019. Linking structural and effective brain connectivity: structurally informed parametric empirical bayes (si-peb). Brain Structure and Function 224, 205–217. doi:10.1007/S00429-018-1760-8/FIGURES/4.

Sokolov, A.A., Zeidman, P., Razi, A., Erb, M., Ryvlin, P., Pavlova, M.A., Friston, K.J., 2020. Asymmetric high-order anatomical brain connectivity sculpts effective connectivity. Network Neuroscience 4. doi:10.1162/netn_a_00150.

Stephan, K.E., Tittgemeyer, M., Knösche, T.R., Moran, R.J., Friston, K.J., 2009. Tractography-based priors for dynamic causal models. NeuroImage 47, 1628–1638. doi:10.1016/J.NEUROIMAGE.2009.05.096.

Stiso, J., Bassett, D.S., 2018. Spatial embedding imposes constraints on neuronal network architectures. Trends in cognitive sciences 22, 1127–1142. doi:10.1016/J.TICS.2018.09.007.

Suárez, L.E., Markello, R.D., Betzel, R.F., Misic, B., 2020. Linking structure and function in macroscale brain networks. Trends in Cognitive Sciences 24, 302–315. doi:10.1016/J.TICS.2020.01.008.

Vázquez-Rodríguez, B., Suárez, L.E., Markello, R.D., Shafiei, G., Paquola, C., Hagmann, P., Heuvel, M.P.V.D., Bernhardt, B.C., Spreng, R.N., Misic, B., 2019. Gradients of structure–function tethering across neocortex. Proceedings of the National Academy of Sciences of the United States of America 116, 21219–21227. doi:10.1073/PNAS.1903403116.

Whitesell, J.D., Liska, A., Coletta, L., Hirokawa, K.E., Bohn, P., Williford, A., Groblewski, P.A., Graddis, N., Kuan, L., Knox, J.E., Ho, A., Wakeman, W., Nicovich, P.R., Nguyen, T.N., van Velthoven, C.T., Garren, E., Fong, O., Naeemi, M., Henry, A.M., Dee, N., Smith, K.A., Levi, B., Feng, D., Ng, L., Tasic, B., Zeng, H., Mihalas, S., Gozzi, A., Harris, J.A., 2021. Regional, layer, and cell-type-specific connectivity of the mouse default mode network. Neuron 109, 545–559.e8. doi:10.1016/J.NEURON.2020.11.011.

